# Capsaicin pretreatment alleviates postoperative pain and reduces primary sensory neuron Ca^2+^ activity

**DOI:** 10.1101/2021.05.21.445191

**Authors:** Hirotake Ishida, Yan Zhang, Ruben Gomez, John Shannonhouse, Hyeonwi Son, Yu Shin Kim

**Affiliations:** Department of Oral & Maxillofacial surgery, University of Texas Health Science Center at San Antonio, 7703 Floyd Curl Drive, San Antonio, TX 78229, U.S.A.; Programs in Integrated Biomedical Sciences, Translational Sciences, Biomedical Engineering, Radiological Sciences, University of Texas Health Science Center at San Antonio, 7703 Floyd Curl Drive, San Antonio, TX 78229, U.S.A.

## Abstract

After surgeries, especially thoracotomy incision, patients develop unbearable pain. Opioids are used for reducing pain but often cause serious side effects. Previously, we found that capsaicin pretreatment of the incision area alleviated spontaneous and thermal pain in a postoperative pain animal model. In the present study, we aimed to monitor primary sensory neuron Ca^2+^ activity in *in vivo* dorsal root ganglia (DRG) in a postoperative pain model using Pirt-GCaMP3 treated with capsaicin or controls. Intraplantar injection of capsaicin (0.05%) alleviated spontaneous, mechanical, and thermal postoperative pain. The Ca^2+^ response in *in vivo* DRG and in *in situ* spinal cord was significantly enhanced in the ipsilateral side compared to contralateral side or naive control. Primary sensory nerve fiber length was significantly decreased in the incision skin area in capsaicin-pretreated animals detected by immunohistochemistry and placental alkaline phosphatase (PLAP) staining. Thus, capsaicin pretreatment alleviates postoperative pain by suppressing Ca^2+^ response due to degeneration of primary sensory nerve fibers in the skin.

## Introduction

Despite many available analgesics, postoperative pain after surgery remains a significant problem and challenge, especially for thoracotomy incision. Around 80% of patients develop serious acute postoperative pain (Apfelbaum et al., 2003). The patients develop spontaneous, mechanical, and thermal pain (Kawamata et al., 2002; Ahmad et al., 2014). Postoperative pain typically lasts over a month after the surgery in up to 50% of patients, and acute postoperative pain sometimes makes a transition to severe chronic pain in up to 10% of patients (Lovich-Sapola et al., 2015). Opioids are one of most common treatment options for postoperative pain. However, serious side effects and consequential opioid addiction problems (Carroll et al., 2012) diminish quality of life of patients (Benyamin et al., 2008). From 5 to 85% of postoperative pain patients develop persistent postoperative pain depending on the surgeries (Macrae, 2008; Thapa and Euasobhon, 2018). Thus, it is important to elucidate the molecular and cellular mechanism underlying postoperative pain to develop more effective drug targets and therapeutics.

Our previous research indicated that intraplantar capsaicin pretreatment before incision alleviates spontaneous and thermal pain due to degeneration of CGRP and IB4/PGP9.5 positive nociceptive nerve fibers in the skin of rats (Kang et al., 2010). In addition, in our experiment with intraplantar capsaicin pretreatment, degeneration of TRPV1-positive primary sensory nerve fibers caused by the capsaicin alleviated spontaneous and thermal pain after plantar incision (Uhelski et al., 2020). Nerve growth factor (Thalakoti, #100) expression level in glabrous skin but not in L4-L6 dorsal root ganglia (DRG) was increased, and increased mRNA levels of insulin-like growth factor-2 (IGF-2) were found using total mRNA sequencing in a rat postoperative pain model (Banik et al., 2005; Tran et al., 2020).

Despite of these attractive findings and studies, *in vivo* dorsal root ganglia activity at the populational level is unclear, and further study might give us a clue to better determine candidate molecules for targeting postoperative pain. Previous studies suggested that enhanced primary sensory neuronal Ca^2+^ signals contribute to postoperative pain (Wang et al., 2000; Joksimovic et al., 2018). However, due to limited evidence from *in vivo* Ca^2+^ imaging, molecular and cellular mechanisms related to Ca^2+^ signals in primary sensory neurons for postoperative pain remain in dispute. We developed populational level, *in vivo* primary sensory neuronal cell body Ca^2+^ imaging using our Pirt-GCaMP3 animal model, in which Ca^2+^ activities of >1,800 neurons in the DRG can be visualized (Kim et al., 2016). Furthermore, Pirt-GCaMP3 enables us to monitor primary sensory nerve fibers in the isolated skin despite being surrounded by many other cell types (Kim et al., 2014). This model permits us to monitor and study Ca^2+^movement in almost all primary sensory neuron cell bodies and fibers using our Pirt-GCaMP3 animals. In the present study, we have assessed *in vivo* primary sensory neuronal cell body Ca^2+^ activity in DRG on a populational ensemble level, and *in situ* Ca^2+^ activity in primary sensory nerve fibers in sliced spinal cord in our postoperative pain model.

## Materials and Methods

### Animals

Mice (C57BL/6J, Jackson Laboratory, Bar Harbor, Maine) were housed in a regular light-dark cycle, with food and water available *ad libitum*. The animal use protocol was approved by the Institutional Animal Care and Use Committee and according to the guidelines for animal experiments approved by the Institutional Animal Care and Use Committee of the University of Texas Health Science Center at San Antonio. Male and female mice were used for all experiments.

### Postoperative pain model

One week after intraplantar injection of capsaicin or vehicle, a 5 mm longitudinal incision was made with a number 11 blade through the skin, fascia, and muscle of right hindpaw under conditions of anesthesia with 2-3% isoflurane. The incision was closed with two single sutures of 6-0 nylon and triple antibiotic ointment was applied.

### Drug treatment

Capsaicin (Sigma, St. Louis, MO) was dissolved in vehicle (1.5% Tween 80, 1.5% ethanol, and 97% saline). Fifty µl of 0.05% capsaicin (100 μg/200 μl) or an equal volume of vehicle was subcutaneously injected to the incision site of the right hindpaw using a 0.5-ml insulin syringe with a 28-gauge needle under conditions of anesthetization using ketamine hydrochloride (80 mg/ml) /xylazine hydrochloride (12 mg/ml) solution, 10 µl/10 g weight. Capsaicin was subcutaneously injected into the intraplantar area 7 days prior to incision surgery.

### Measurement of 50% paw withdrawal threshold

von Frey filament was applied to plantar surface and escape behavior was observed. A mouse was placed in a transparent plastic cage on a metal grid and allowed to habituate for at least 1 hour before testing. von Frey filament intensity from 0.04 (No. 2.44) to 2.0□g (No.4.31) was used with an up-down test. The filament was bent for 3 seconds after contacting to plantar skin. The interval between each application was 3 min after escape behavior was observed, or 30 seconds if escape behavior was not observed. To calculate the 50% threshold of paw withdrawal, the following formula was used: 10^(xf+kx0.22)^/10,000. X_f_ =□the value of the final filament used (in log units), k□=□a value based on the response pattern, as reported by Chaplan et al.,(Chaplan et al., 1994).

### Measurement of thermal pain test

To measure the thermal response, a Hargreaves test was performed. A mouse in a transparent plastic cage on heated glass board at 30°C was allowed to habituate for at least 1 hour before testing. Radiant heat was applied to the mouse plantar area through the glass, and the duration until escape behavior was measured. Initial heat applied was 34°C, which increased to 51°C in 10 seconds. This process was done 5 times, and the average time to escape behavior was calculated. The interval between each trial was at least 5 minutes.

### Measurement of spontaneous foot lifting

Spontaneous foot lifting was used as a measure of spontaneous pain. A mouse in a transparent plastic cage on a metal grid was allowed to habituate for at least 1 hour before testing. An angled mirror was set under the metal grid during habituation and mouse behavior reflected by the mirror was recorded with video camera for 10 minutes. Frequency of spontaneous foot lifting and foot licking of the incision foot was determined.

### Open field test

Three hours after incision surgery, an open field test was performed. To monitor mouse behavior, a mouse was put in a new cage surrounded by white square box. The distance between the white board and side of the cage was around 10 cm. Behaviors were recorded for 10 min using a camera mounted above the cage, and were assessed as total walking distance. Movement was calculated using Image J software with the Animal Tracker plugin (Gulyás et al., 2016).

### *In vivo* DRG Ca^2+^ imaging

One day after incision surgery, *in vivo* Ca^2+^ imaging in the DRG was performed. L5 whole DRG Ca^2+^ imaging in a live mouse was performed for 2-4 hour after DRG exposure as we previously described (Kim et al., 2016). Rectal temperature was monitored as body temperature and kept at around 37°C using a heat pad. Mouse movement due to breath and heart beat was minimized using clamps on vertebra bone. The mouse was anesthetized with ketamine hydrochloride (80 mg/ml) /xylazine hydrochloride (12 mg/ml) solution mixed with saline (1:1), 10 µl/10 g weight. Live images were obtained using single photon confocal microscopy (Carl Zeiss, Jena, Germany) from 10 frames at 465-960 ms/frame in each z-axis (512 × 512 or 1024 × 1024 pixels in the x-y plane) using a 10X dry objective lens. Solid diode lasers at 488 and 532 nm wavelength were used for emission at 500-550 nm for green fluorescence and 550-650 nm for red fluorescence, respectively. The depth of each x-y focal plane where the laser captured images was from 0 to about 100 µm. Fifteen image cycles were obtained, and the duration time of each cycle was around 4.5 to 7.5 seconds. Small brush, large brush, 0.4 g, and 2.0 g von Frey filament were applied to the hindpaw of exposed DRG side. A press of 100 g and 300 g were applied to the whole palm of hindpaw using a rodent pincher (Bioseb, U.S.A.). Whole hindpaw was immersed into 50°C water, and acetone was applied by pipette to the hindpaw. One hundred mM KCl (30 μl) was subcutaneously injected into the hindpaw using a 0.5-ml insulin syringe with a 28-gauge needle. Small and large brush bristles were 5 mm and 40 mm, respectively.

### *In Situ* Ca^2+^ imaging in spinal cord slice

Animals were killed by decapitation under conditions of anesthesia using ketamine. The lumbar enlargement area (L3-5) was removed, and placed in ice-cold NMDG solution (N-methyl-D-glucamine 135 mM, KCl 1 mM, MgCl_2_ 1.5 mM, KH_2_PO_4_ 1.2 mM, CaCl_2_ · 2H_2_O 0.5, C_5_H_14_NO · HCO_3_ 29.5 mM, glucose 10.7 mM) aerated with (5% CO_2_, 95%O_2_). Spinal cord was dissected and transversely sliced at 300-400 µm using a Leica VT1200S vibratome at 4°C. The spinal cord slices were immediately placed in synthetic interstitial fluid (NaCl_2_ 107.8 mM, KCl 3.5 mM, MgSO_4_ · 7H_2_O 0.69, NaH_2_PO_4_ · 2H_2_O 1.67 mM, glucose 5.55 mM, sucrose 7.6 mM, CaCl_2_ · 2H_2_O 1.53 mM, NaHCO_3_ 26.2 mM, gluconic acid sodium salt 9.64 mM) aerated with (5% CO_2_, 95%O_2_) on an imaging stage (Siskiyou Corporation, Grants Pass, OR, U.S.A.) stabilized with Harp (Scientific Instruments, Farmingdale, NY, U.S.A.). The dorsal horn of the tissue was viewed using a 40X water immersion objective lens (Carl Zeiss, Jena, Germany). The z-axis of spinal cord was imaged at a depth of 100 to 150 μm at approximately 10 µm intervals. A solid diode laser at 488 nm was used for emission at 500-550 nm for green fluorescence. During Ca^2+^ imaging experiments at room temperature, synthetic interstitial fluid solution aerated with 95% O_2_/5% CO_2_ was perfused at 400 ml/hour. 2-D images of primary sensory nerve fiber were made from merged z-axis maximum fluorescence intensities (512 × 512 pixels) (Kim et al., 2014). Green fluorescence from GCaMP3 was analyzed by image J (NIH, U.S.A.).

### Ca^2+^ transient calculation

Ca^2+^ transients in *in vivo* DRG and in spinal cord slice imaging were determined using the formula: ΔF/ F_0_, where F_0_ represents basal intensity at 500-550 nm for green fluorescence, and ΔF represents intensity calculated by subtracting basal intensity from the intensity at each time point.

### Cell size measurement

Cell size was defined by the average of the long diameter and short diameter from one cell body using Zen Blue software (ZEN3.1, Jena, Germany).

### Immunohistochemistry

Plantar skin was dissected one day after surgery, and washed in 0.1 M phosphate buffered saline (PBS) followed by 4% paraformaldehyde (PFA) overnight at 4°C. Tissue was transferred to 10% sucrose overnight at 4°C followed by 30% sucrose, and frozen at −80°C. Frozen skin was returned to room temperature, transferred to optimal cutting temperature (OCT) embedding medium (Fisher Healthcare), and frozen at −20°C. Tissue was sliced (20 µm) using a cryostat. Sections were transferred to slides and dried at 37°C for an hour, incubated with guinea pig anti-TRPV1 antibody (1:250, cat. no. ab10295, Abcam, Cambridge, MA, USA), chicken anti-GFP antibody (1:250, cat. no. CH23105, Neuromics, Edina, MN, USA), and rabbit anti-β-III tubulin antibody (1:250, cat. no. ab18207, Abcam) overnight at 4°C incubation of blocking solution. Primary antibody was washed out with PBS, and slides were incubated for 90 min at room temperature with Goat secondary antibody to guinea pig IgG - Alexa Fluor 568 (1:200, Invitrogen, Waltham, MA, USA), goat secondary antibody to Rabbit IgG - H&L (Cy5) (1:200, cat. no. A-21244) and goat secondary antibody to chicken IgG Alexa Fluor 488 (1:200, Invitrogen). Sections were washed three times in PBS, and a cover glass was applied with Prolong™ Diamond Antifade Mountant with DAPI (Invitrogen). Immunofluorescence was viewed using a 10X dry or 40X water immersion objective lens by single photon confocal microscopy (Carl Zeiss, Jena, Germany). Human placental alkaline phosphatase (PLAP) is a useful protein to trace peripheral axons in dissected tissue and whole mount skin. We purchased PLAP floxed mice from Jackson Lab (Bar Harbor, ME), and used the experimental method described by Dr. Jeremy Nathans at Johns Hopkins University (Chang et al., 2014). After 4% PFA fixation, skin tissue was fixed on sylgard in a 10 cm glass dish, and then transferred into PBS containing 1 mM MgCl_2_. The dish was gently rotated in a water bath at 70°C for 90 min. Skin tissue was transferred to BCIP/NBT membrane alkaline phosphatase substrate solution (Rockland), and incubated around 36 hours at room temperature. To stop the reaction, skin tissue was washed tree times with PBS/0.1% tween-20 over 1 hour. After washing, skin tissue was moved to ethanol to dehydrate, and then pinned on sylgard with benzyl benzoate and benzyl alcohol (BBBA) for 3 hours. Skin tissue was placed between glass plates, and any bubbles and extra BBBA were removed. PLAP staining was imaged using a bright-field microscope with 10X and 40X objectives.

### Drugs

Capsaicin was purchased from Sigma-Aldrich (St. Louis, MO, USA), and wortmannin was purchased from AdooQ science (Irvine, CA, U.S.A); other drugs were purchased from Fisher bio-reagents (Pittsburgh, PA, U.S.A).

### Statistical analysis

Results are expressed as mean□±□S.E.M. Student’s *t*-test or two-way ANOVA comparison test was used to analyze differences. For multiple comparisons, Dunnett’s multiple test was performed. *P*□<□0.05 was considered statistically significant.

## Results

### Subcutaneous intraplantar injection of capsaicin alleviates spontaneous, mechanical, and thermal pain evoked by intraplantar incision

An incision of the plantar area induced mechanical and thermal pain that was assessed at 3 hours, 1 day, 2 days, and 3 days after the surgery (Figure 1A-1D). The injection of capsaicin 7 days prior to the incision surgery alleviated the mechanical and thermal pain induced by the incision surgery (Figure 1E & F). The injection of capsaicin also decreased spontaneous foot lifting following incision surgery (Figure 2A). In open field tests, total walking distance and average walking velocity from incised mice were reduced compared to naïve and capsaicin-treated mice following the incision (Cap-ICS mice) (Figure 2B-D).

**Figure 1.**
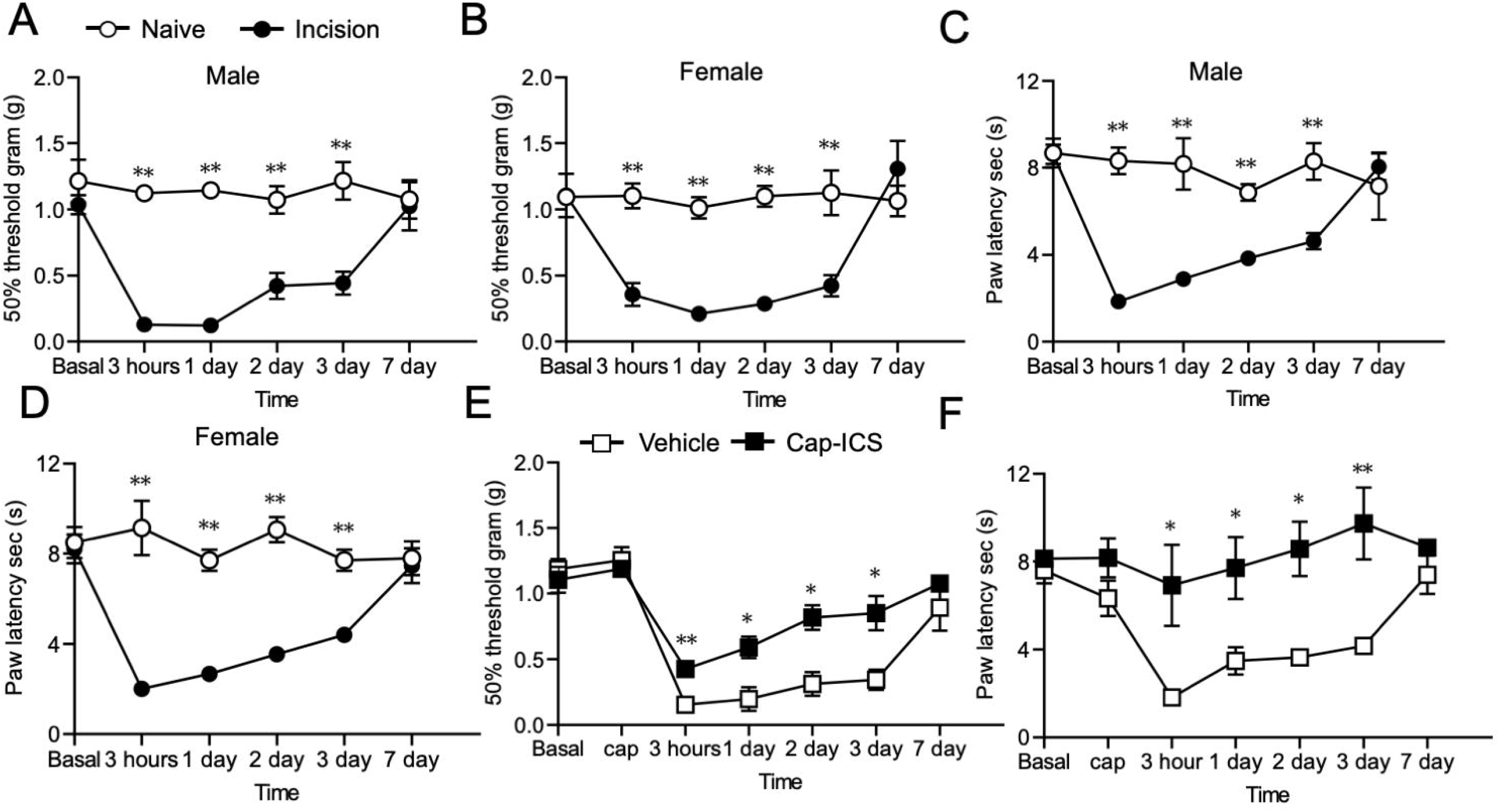
Capsaicin pretreatment (intraplantar injection, 0.05% capsaicin) significantly alleviates postoperative mechanical and thermal pain in a mouse model. **A, B**. Mechanical pain test using von Frey filament was performed after incision surgery at 3 h, 1 d, 2 d, 3 d, and 7 d in male and female mice. Mechanical sensitivity is plotted as 50% withdraw threshold in grams. **C, D**. Thermal pain test using Hargreaves method was performed after incision surgery at 3 h, 1 d, 2 d, 3 d, and 7 d in male and female mice. Thermal sensitivity is plotted as paw withdraw latency following thermal stimulus. **E, F**. Mechanical or thermal pain test using von Frey filament or Hargreaves method was performed after incision surgery at 3 h, 1 d, 2 d, 3 d, and 7 d in vehicle or capsaicin-treated (7 days prior to incision surgery) incision mouse model. Cap-ICS: capsaicin-treated incision. Fig. A and C, naïve: n = 5, incision: n = 7; Fig. B and D, naïve: n = 5, incision: n = 6; Fig. E and F, vehicle: n =5, Cap-ICS: n = 5. Data represent mean ± S.E.M.; T-test, \**p*<0.05, \*\**p*<0.01 vs. naïve or vehicle.

**Figure 2.**
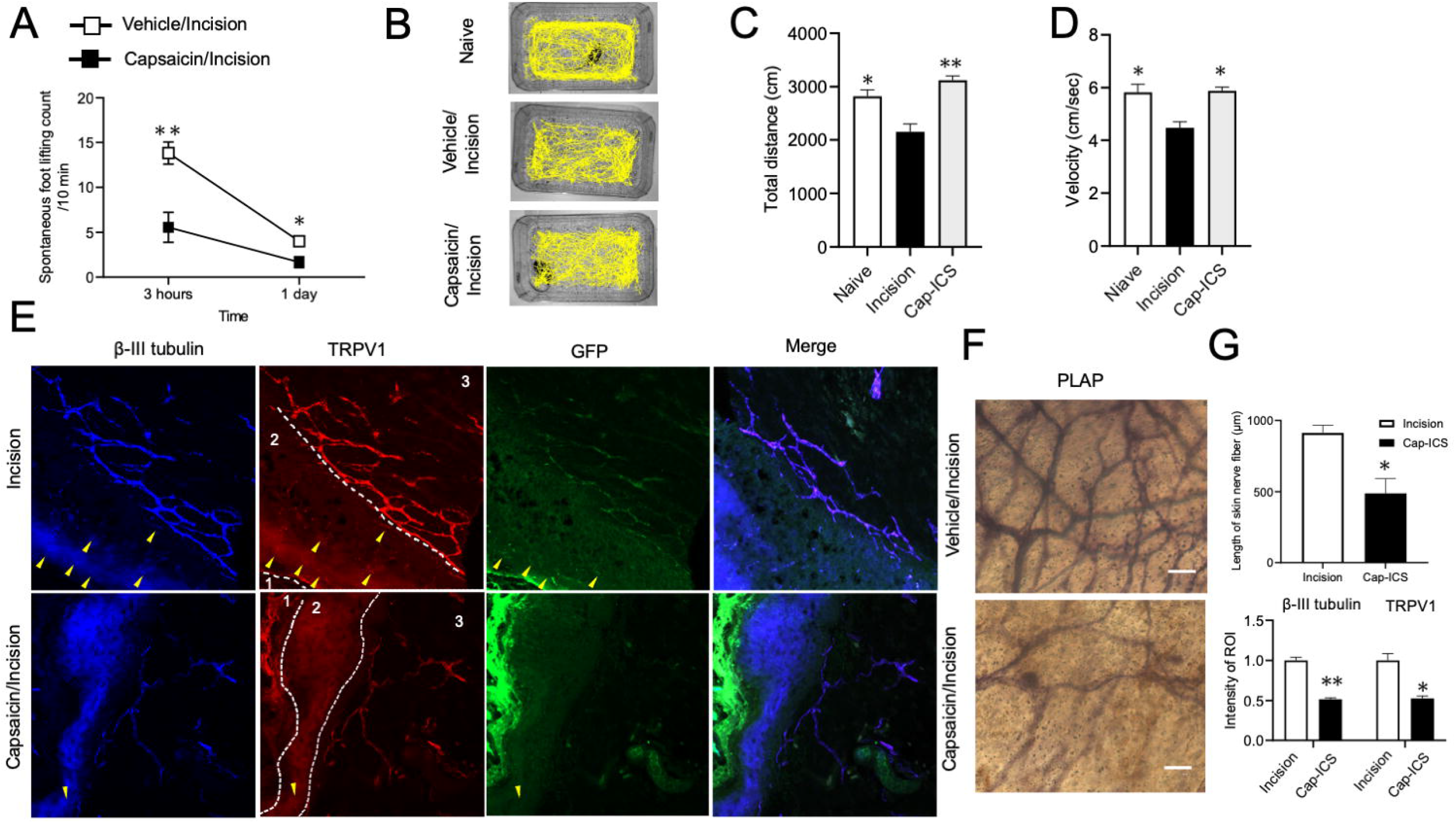
Capsaicin pretreatment (intraplantar injection, 0.05% capsaicin) significantly alleviates spontaneous pain and decreases the length of peripheral primary sensory nerve fiber in the skin. **A**. Spontaneous foot lifting as an index of spontaneous pain was counted at 3 h and 1 d after incision surgery. The number of spontaneous foot lifts by vehicle and capsaicin-treated (7 days prior to incision surgery) mice is plotted. **B**. Examples of traces of mouse movement in the open field test for assessment of spontaneous pain in naïve, vehicle-treated mice, and capsaicin-treated mice. **C**. Total walking distance in open field test of naïve, vehicle-treated, and capsaicin-treated mice. **D**. Average walking velocity in the open field test of naïve, vehicle-treated, and capsaicin-treated mice. **E**. Confocal microscopic immunohistochemistry image (40X) of β-III tubulin (sensory nerve fiber), TRPV1 channel, and GCaMP (GFP)-expressing primary sensory neuron fibers in skin from incision and capsaicin-treated Pirt-GCaMP3 mice. Area 1, skin surface; area 2, epidermis; area 3, dermis. Arrowheads indicate nerve fibers and nerve endings in epidermis. **F**. PLAP staining image (40X) in skin from vehicle-treated and capsaicin-treated Pirt-cre/PLAP^flox^ mice. Black lines are sensory nerve fibers. **G**. Graph of PLAP-positive skin nerve fiber length and ratio of fluorescence intensities of β-III tubulin and TRPV1 channels from either incision only mice or capsaicin-treated incision mice. Cap-ICS, capsaicin-treated incision. Scale bars, 100 µm. Fig. A, incision, n = 6; Cap-ICS, n = 9. Fig. B-D, naïve, n = 5; incision, n = 5; Cap-ICS, n = 5. Fig. E and G, incision, n = 3; Cap-ICS, n = 3. Fig. F and G, incision, n = 3; Cap-ICS, n = 3. Data represent mean ± S.E.M.; T-test or Dunnett’s test; \**p*<0.05, \*\**p*<0.01 incision vs. naïve or capsaicin-treated incision.

### The length and intensity of TRPV1 and placental alkaline phosphate (PLAP)-positive primary sensory nerve fibers are decreased in the capsaicin-treated incision model

We performed intraplantar injection of capsaicin to induce degeneration of TRPV1-positive nociceptive primary sensory nerve fibers in plantar skin. To check the effect of capsaicin injection, expression of transient receptor potential vanilloid (TRPV1) channel-positive primary sensory nerve fibers were measured by IHC. TRPV1-positive primary sensory nerve fibers in the incision skin were decreased by the intraplantar capsaicin injection compared to control incisions (Fig 2E & G). Furthermore, β-III tubulin-positive skin nerve fibers in capsaicin-treated mice were decreased compared to untreated mice in the incision model (Fig 2E & G). Strong PLAP expression in primary sensory nerve fibers are detected in almost all primary sensory nerve fibers in the skin (Wu et al., 2012). We tried to visualize PLAP expression driven by Pirt-cre in all primary sensory nerve fibers using skin flat mount PLAP staining. Primary sensory nerve fiber lengths in capsaicin-treated incision model were dramatically decreased compared to vehicle-treated incision mice (Fig 2 F & G). Morphological changes in primary sensory nerve fibers in capsaicin-treated incision mice were detected, showing weak staining and fewer branching fibers than incised controls (Fig 2 F & G).

### Plantar incision causes spontaneous Ca^2+^ activity which is dramatically reduced by injection of capsaicin *in vivo* in Pirt-GCaMP3 mice

To elucidate the relationship between spontaneous pain and primary sensory neuron Ca^2+^ movement in DRG, we performed primary sensory neuron soma Ca^2+^ imaging 1 day after the incision surgery. As in our previous study, using single photon confocal microscopy, we monitored Ca^2+^ activity and Ca^2+^ transient response of primary sensory neuron soma in Pirt-GCaMP3 mice with mechanical and thermal stimulation of the paw (Fig 3). We confirmed that GCaMP3 is expressed in cytosol, shown by the lack of signal in the nucleus, seen as the unstained area in the soma center (Fig 3A). In this experiment, we define *in vivo* “spontaneous Ca^2+^ activity” as occurring under conditions of no stimulation to the paw or to the animal. Spontaneous Ca^2+^ activity and Ca^2+^ oscillation events were detected in a larger number of cell soma in incision mice than in naïve mice (Fig 3B-D). The increased Ca^2+^ activity of primary sensory soma was reduced in Capsaicin-treated incision mice (Fig 3B). Spontaneous Ca^2+^ activity in DRG neuronal cell bodies did not synchronize and were random events (Fig 3C).

**Figure 3.**
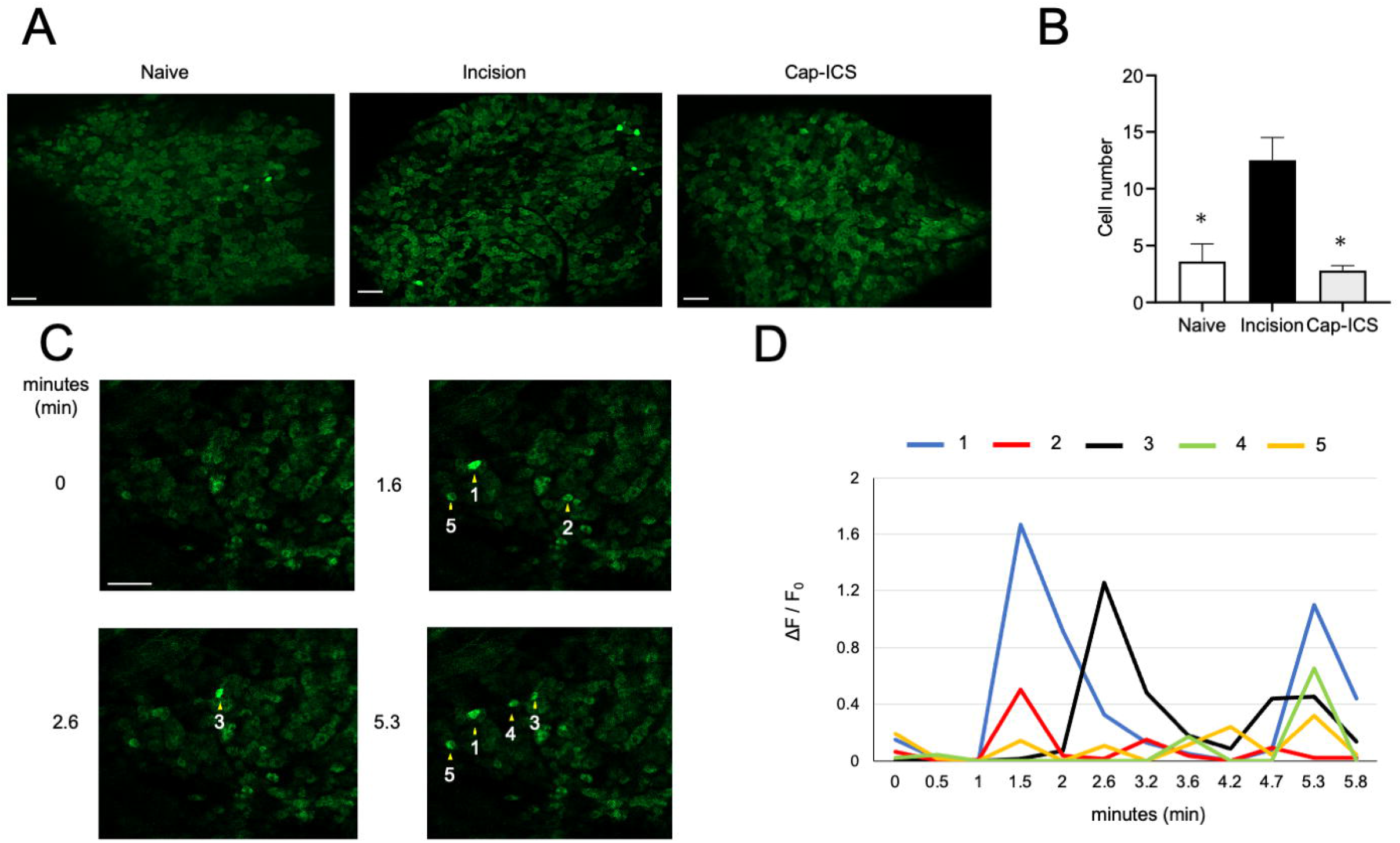
Spontaneous Ca^2+^ activity of primary sensory neurons in *in vivo* entire dorsal root ganglia (DRG) imaging using Pirt-GCaMP3 mice. **A**. Representative high resolution images of DRG green fluorescence derived from GCaMP3 in naïve, incision, and capsaicin-treated incision mice. **B**. Total cell numbers exhibiting spontaneous Ca^2+^ activity in naïve, incision, and capsaicin-treated incision mice. **C**. Representative spontaneous Ca^2+^ transient responses of primary sensory neurons in incision model. Each arrowhead indicates a spontaneous Ca^2+^ transient-positive cell. **D**. Ca^2+^ transients in spontaneous activity-positive cells of Figure 3C. Cap-ICS, capsaicin-treated incision. Scale bars, 100 µm. Naïve, n = 5; incision, n = 4; Cap-ICS, n = 5. Data represent mean ± S.E.M.; Dunnett’s test; \**p*<0.05, \*\**p*<0.01, incision vs. naive or capsaicin-treated incision.

### Activation of many primary sensory neurons in the incision model were detected by weak stimulation to the hindpaw plantar skin

By the von Frey filament test, mechanical threshold in the incision animal model was much lower than in naïve or capsaicin-treated incision animals (Fig 1). We examined Ca^2+^ response induced by small & large brush, and by 0.4 g & 2.0 g von Frey filament application. Many more cells were activated by small & large brush in the incision model than in naïve or capsaicin-treated animals, but this was not seen following 0.4 g & 2.0 g von Frey filament stimuli (Fig 4 A, B). No significant differences in Ca^2+^ transients induced by large brush stimulation were detected among groups (naïve, incision, capsaicin-treated incision) (Fig 4C).

**Figure 4.**
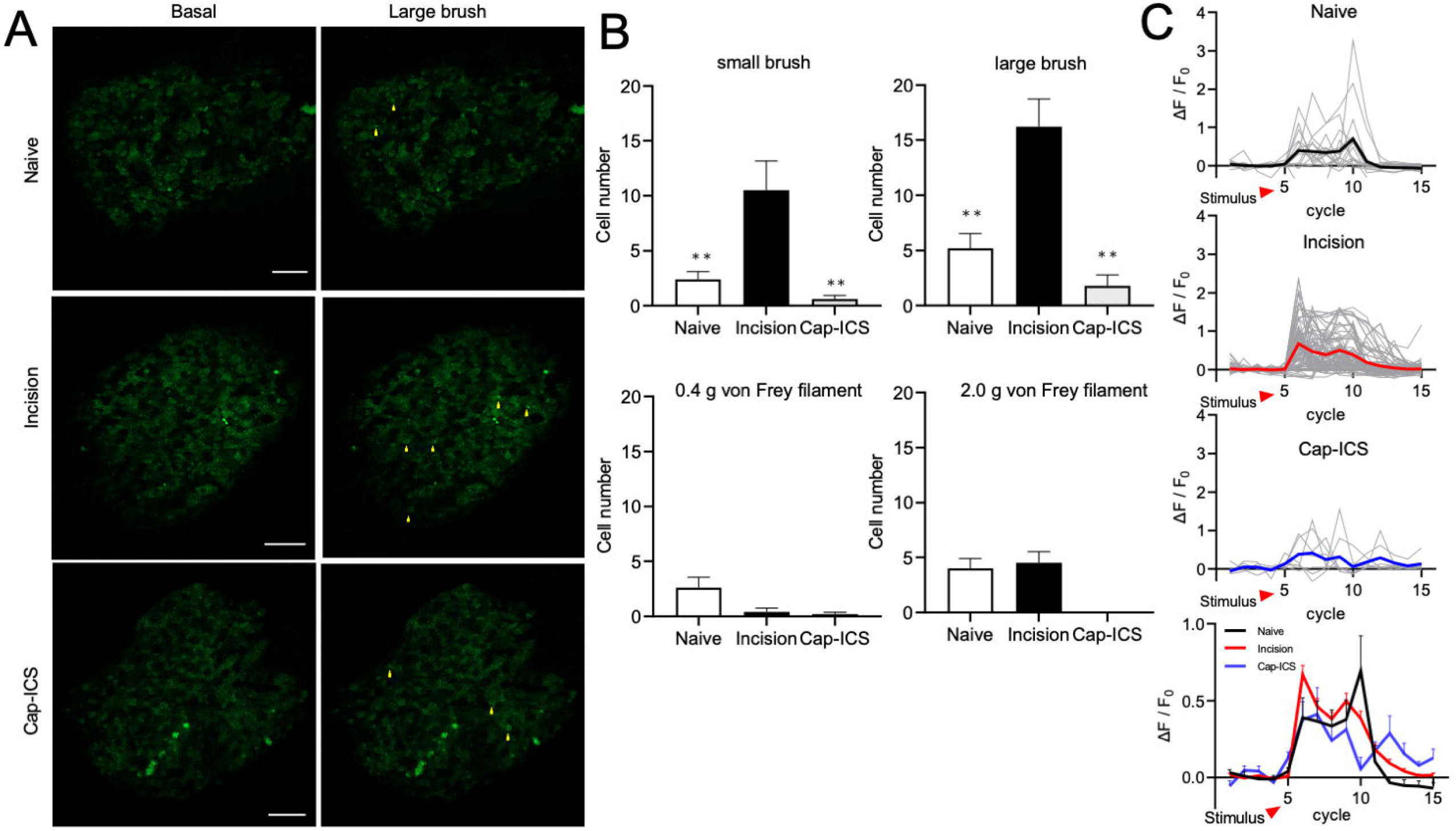
*In vivo* DRG imaging of Ca^2+^ activity induced by weak mechanical stimulation of primary sensory neurons. **A**. Representative Ca^2+^ elevation images induced by large brush stimulation in naïve, incision, and capsaicin-treated incision mice. **B**. Total cell number activated by stimulation with small brush, large brush, 0.4 g von Frey, and 2.0 g von Frey filament in naïve, incision, and capsaicin-treated incision mice. **C**. Ca^2+^ transient induced by large brush stimulation in naïve, incision, and capsaicin-treated incision mice. Black (naïve mice), red (incision mice), and blue (capsaicin-treated incision mice) lines indicate average Ca^2+^ transient. Cap-ICS, capsaicin-treated incision. Scale bars, 100 µm. Naïve, n = 5; incision, n = 4-5; Cap-ICS, n = 5. Data represent mean ± S.E.M. Dunnett’s test; \**p*<0.05, \*\**p*<0.01, incision vs. naïve or capsaicin-treated incision mice.

### A larger number of cells in primary sensory neurons are activated by mechanical, thermal and chemical stimuli to the hindpaw

A larger number of cells were activated by 100 g press, 300 g press, 50°C thermal, or 100 mM KCl intraplantar injection in incision animals compared to naïve or capsaicin-treated incision animals (Fig 5 A & B). However, no significant differences in Ca^2+^ activated cell numbers from acetone application were detected between groups (naïve, incision, capsaicin-treated incision) (Fig 5B). Ca^2+^ transients in the incision model induced by 100 g press, 50°C thermal, or 100 mM KCl intraplantar injection were greatly enhanced compared to naïve or capsaicin-treated incision animals (Fig 5C).

**Figure 5.**
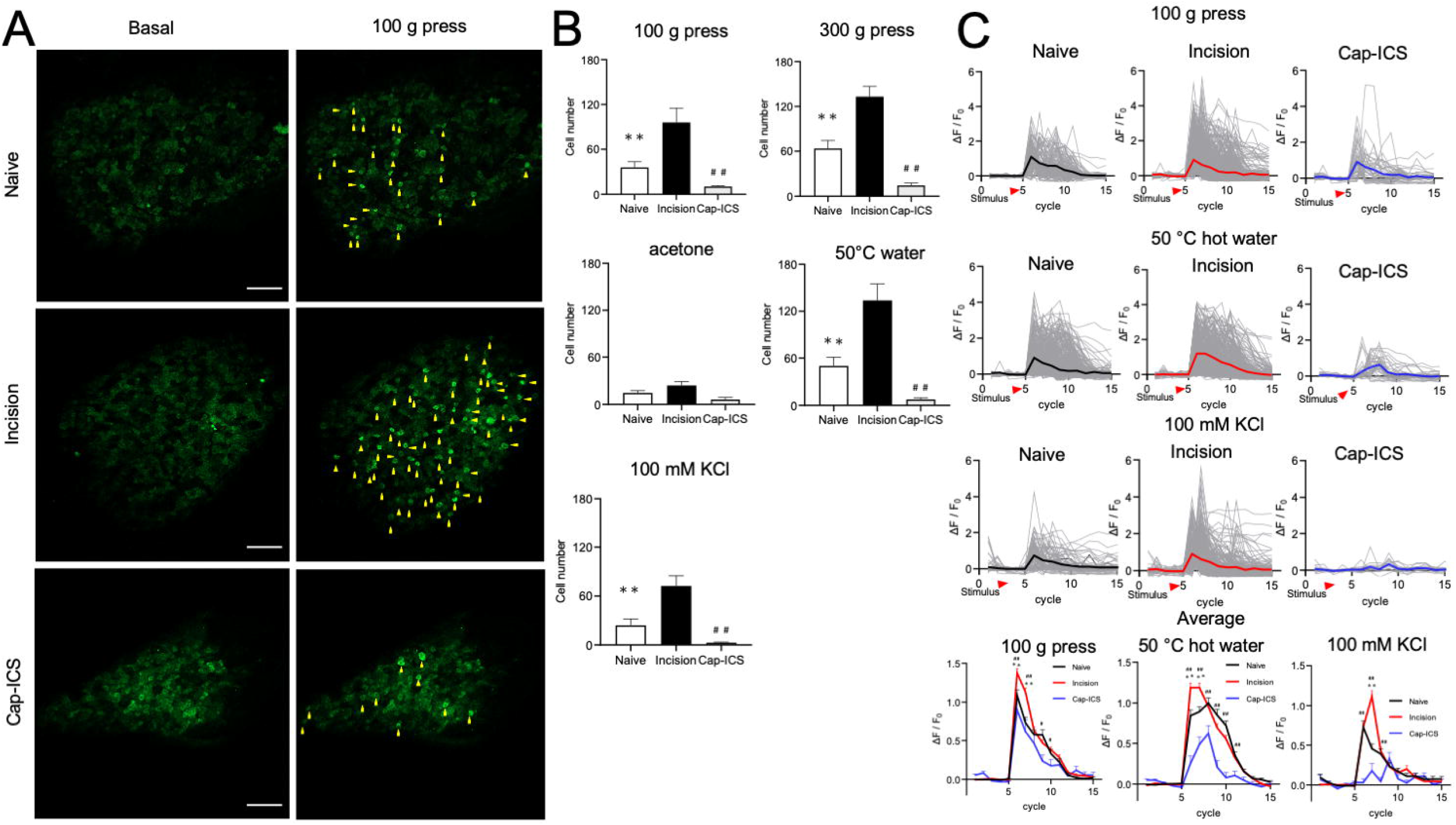
*In vivo* DRG imaging of Ca^2+^ elevation induced by medium to strong stimuli in primary sensory neurons. **A**. Representative Ca^2+^ elevation images induced by 100 g press to the hindpaw in naïve, incision, and capsaicin-treated incision mice. **B**. Total cell numbers activated by 100 g press, 300 g press, or hot water to the hindpaw, or acetone to the plantar area, or 100 mM KCl intraplantar injection in naïve, incision, and capsaicin-treated incision mice. **C**. Ca^2+^ transient induced by 100 g press or 50° hot water to the hindpaw, or 100 mM KCl intraplantar injection in naïve, incision, and capsaicin-treated incision mice. Black (naïve mice), red (incision mice), blue (capsaicin-treated incision mice) lines indicate average Ca^2+^ transients. Cap-ICS, capsaicin-treated incision. Scale bars, 100 µm. Naïve, n = 4-5; incision, n = 3-5; Cap-ICS, n = 5. Data represent mean ± S.E.M. Dunnett’s test; \**p*<0.05, \*\**p*<0.01 incision vs. naïve mice; ^#^*p*<0.05, ^##^*p*<0.01 incision vs. capsaicin-treated incision mice.

### A larger number of small and medium diameter cells of primary sensory neurons are activated by stimuli to the hind paw in the incision model

We analyzed diameters of primary sensory neurons activated by the stimuli in naïve, incision, and capsaicin-treated incision. Cell numbers of small diameter neurons showing spontaneous Ca^2+^ activity and cell numbers activated by 100 g press, 50°C, and 100 mM KCl intraplantar injection in incision mice were increased compared to naïve and capsaicin-treated incision mice (Fig 6A). Cell numbers of medium diameter neurons activated by large brush, small brush, 100 g press, 300 g press, 50°C, and 100 mM KCl intraplantar injection were greatly increased in incision mice compared to naïve and capsaicin-treated incision mice (Fig 6B). The numbers of medium diameter neurons activated by small brush, large brush, and 300 g press were increased in incision compared to capsaicin-treated incision mice (Fig 6C).

**Figure 6.**
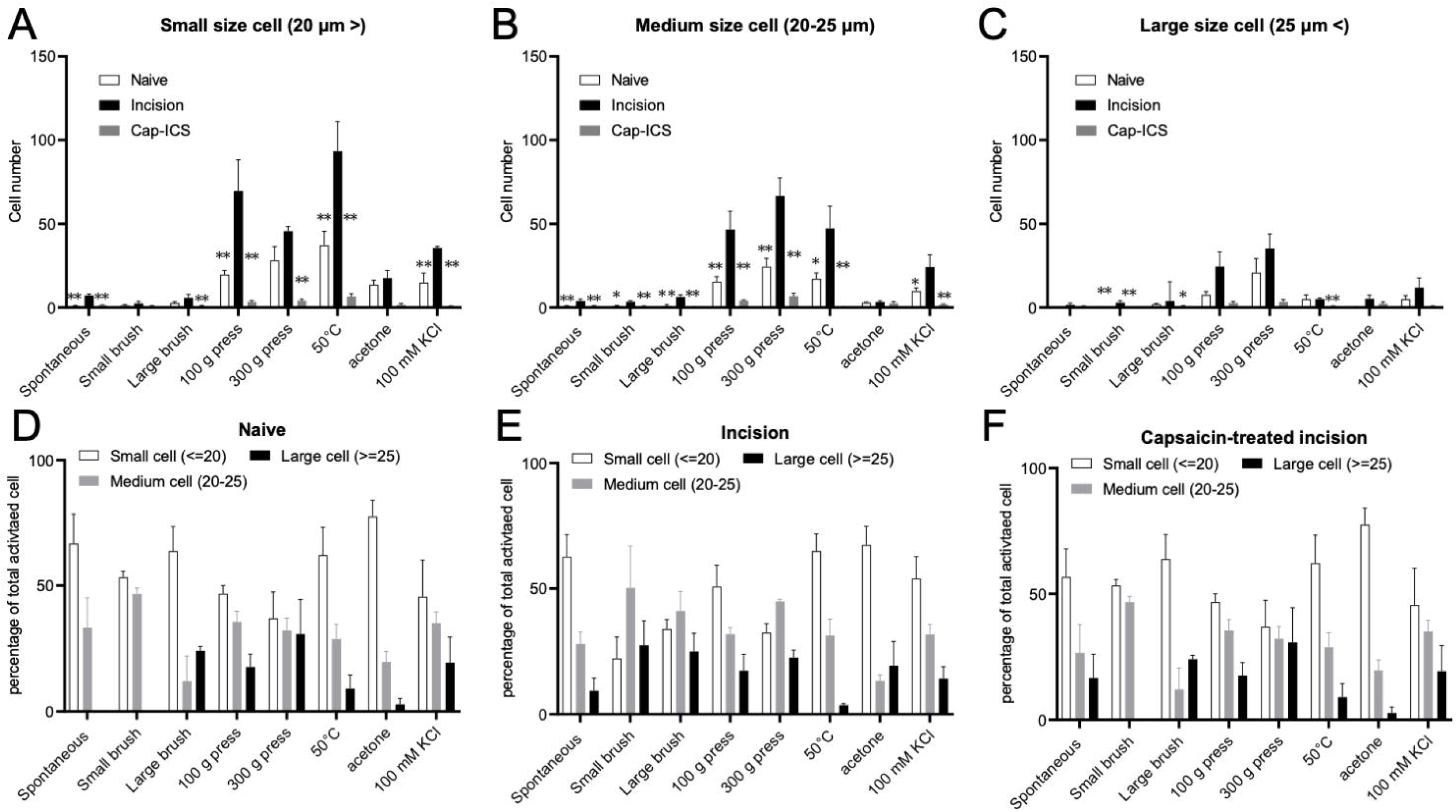
Primary sensory neurons involved in postoperative pain are mainly small and medium diameter. **A-F**. Analysis of cell diameters of primary sensory neurons activated by stimuli to the hindpaw in naïve and incision mice. Cap-ICS, capsaicin-treated incision. Naïve, n = 4-5; incision, n = 3-5; Cap-ICS, n = 5. Data represent mean ± S.E.M. Dunnett’s test; \**p*<0.05, \*\**p*<0.01 incision vs. naïve or capsaicin-treated incision mice.

### *In situ* primary sensory nerve fiber Ca^2+^ imaging in spinal cord sections

We determined that enhanced Ca^2+^ signaling in the incision model is dependent on the activation of capsaicin-sensitive primary sensory neurons. We examined Ca^2+^ transients from primary sensory nerve fibers in lamina I and II in the incision model using spinal cord slices from Pirt-GCaMP3 mice. Ca^2+^ transients induced by application of 1 μM capsaicin or 50 mM KCl to the ipsilateral side of the incision were significantly enhanced compared to applications to the contralateral side (Figure 7 A & C). However, no significant differences in Ca^2+^ transients induced by 100 mM KCl were detected between applications to the contralateral and ipsilateral side of the incision (Figure 7 A). In capsaicin-treated incision mice, no significant differences were detected between Ca^2+^ transients induced by applications of capsaicin (1 µM), 50 mM KCl, or 100 mM KCl to the ipsilateral and contralateral side of the incision (Figure 7 B).

**Figure 7.**
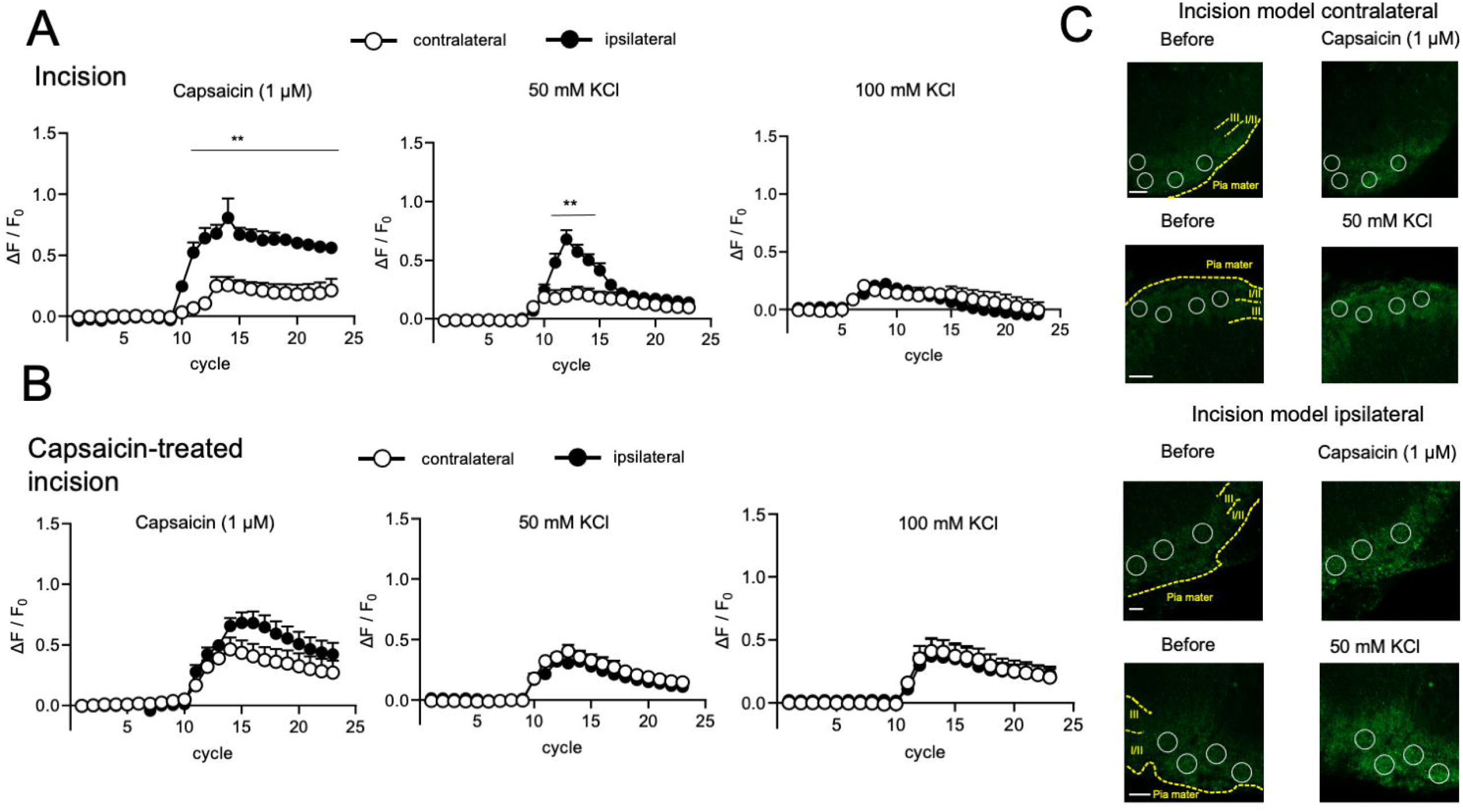
Ca^2+^ transient imaging of primary sensory neuronal central terminals in *in situ* sliced spinal cord in the postoperative pain model using Pirt-GCaMP3 mice. **A & B**. Measurement of Ca^2+^ transients in *in situ* sliced spinal cord from vehicle-treated (A) or capsaicin-treated (B) incision Pirt-GCaMP3 mice. Ca^2+^ transients were induced by bath application of 1 μM capsaicin, 50 mM KCl, or 100 mM KCl. **C**. Representative Ca^2+^ transient images of primary sensory neuronal central terminal induced by capsaicin (1 µM) and 50 mM KCl in ipsilateral and contralateral side of sliced spinal cord from vehicle-treated or capsaicin-treated incision mice. White circles show regions of interest (ROI). Scale bars, 50 µm. Incision mice, n = 4; Capsaicin-treated incision mice, n = 3. Data represent mean ± S.E.M. T-test; \**p*<0.05, \*\**p*<0.01 ipsilateral vs. contralateral.

## Discussion

The present study demonstrates that *in vivo* spontaneous Ca^2+^ activity at cell bodies of primary sensory neurons, and *in vivo* Ca^2+^ responses to mechanical, thermal, and chemical stimuli, are highly enhanced postoperatively. The Ca^2+^ response in central terminals of primary sensory neurons are also significantly enhanced in ipsilateral spinal cord of postoperative pain model. The enhancement of spontaneous Ca^2+^ activity and Ca^2+^ response is diminished due to sensory nerve degeneration induced by capsaicin intraplantar injection prior to surgery. The present study also finds that capsaicin pretreatment alleviates spontaneous, mechanical, and thermal pain by degeneration of skin nerve fibers including TRPV1-positive primary sensory neuron fiber. Our present study suggests potential clinical usefulness of capsaicin pretreatment or nociceptive nerve block treatment prior to surgery to alleviate postoperative pain.

Previously, we showed that both 1% (200 µl) and 0.05% (200 µl) capsaicin intraplantar injection alleviated spontaneous and thermal pain but not mechanical pain, and did not impact wound healing in a rat postoperative pain model (Kang et al., 2010; Uhelski et al., 2020). In the present study, 50 µl of 0.05% capsaicin intraplantar injection alleviated spontaneous, mechanical, and thermal pain in a mouse postoperative pain model. In this mouse postoperative pain model, mast cell number was significantly increased in the surgical site, and was mediated by Mas-related G-protein-coupled receptor in response to substance P released by primary sensory nerve fibers in the inflamed skin, contributing to swelling of skin around the wound followed by mechanical and thermal pain (Green et al., 2019). In addition, TRPV1-positive primary sensory neurons express substance P, and TRPV1 ion channels facilitate substance P release in inflammatory conditions (Hsieh et al., 2012; Li et al., 2019). TRPV1 conditional knock-out alleviated thermal pain but not mechanical pain induced by plantar incision. In contrast, our subcutaneous capsaicin injection significantly alleviated mechanical pain. Because TRPV1 was knocked out ubiquitously from early embryogenesis in the previous study, pain behavior phenotypes evoked by primary sensory neuron would likely be different from pain behaviors in our model in which TRPV1-positive nerve degeneration follows intraplantar capsaicin injection. We confirmed that topical application of 100 µM capsaicin to the DRG elevated the cell body Ca^2+^ level of primary sensory neurons (data not shown). Furthermore, piezo channels are top candidates that might play a major role in mechanical pain. Piezo 1 channels show co-expression with TRPV1 channels in DRG as well (Roh et al., 2020). Roh et al., found that Yoda 1, Piezo 1 channel agonist, and capsaicin induced inward current in around 90% of TRPV1-positive DRG neurons (Roh et al., 2020). TRPV1-positive primary skin nerve degeneration induced by capsaicin pretreatment reduces mechanical pain; as well, Piezo1 is expressed in TRPV1-positive neurons, suggesting a role for Piezo1 in postoperative mechanical pain. It might be possible that species differences between rats and mice in degeneration mechanisms of TRPV1 channel-expressing primary sensory neurons would have an impact on mechanical pain induced by plantar incision.

To assess the effects of intraplantar capsaicin injection onto skin nerve fiber, we visualized primary sensory nerve fibers by immunohistochemistry using placental alkaline phosphatase (PLAP), β-III tubulin, TRPV1 channel, and GFP antibodies. PLAP histochemistry was used to visualize genetically-labeled axon arbors (Wu et al., 2012). Specific expression of PLAP on primary sensory neurons was driven by primary sensory neuron-specific Pirt-cre. β-III tubulin is regarded as a neuronal fiber marker, and is used for detection of primary sensory nerve fiber (Katsetos et al., 1993; Dráberová et al., 1998; Farahani et al., 2019). Around 30%-40% of total primary sensory fibers are TRPV1-positive (Bär et al., 2004; Binzen et al., 2006), and capsaicin pretreatment induced degeneration of almost all TRPV1-positive primary sensory nerve fibers. The present results suggest that degeneration of skin nerve fibers induced by capsaicin can alleviate mechanical and thermal pain caused by activation of TRPV1 channels and TRPV1-positive nerve fibers.

In the present study, we assessed spontaneous pain by counting spontaneous foot lifting and conducting an open field test. Foot lifting events associated with postoperative pain in animals treated with capsaicin were significantly decreased compared to untreated controls. This result suggests that spontaneous Ca^2+^ activity of DRG primary sensory neurons induced by TRPV1-positive nerve fibers plays a key role in generation of spontaneous pain. Mediators in DRG and skin, or changes in peri-environmental primary sensory neuron fibers in the skin, could persistently activate TRPV1-positive nerve fibers through activation of TRPV1 channels. Low pH (4.0-5.0) of local skin caused by incision-mediated inflammation will activate TRPV1 and other channels (Tominaga and Tominaga, 2005; Schreml et al., 2010). We also examined spontaneous pain using an open field test. Animals with plantar incision significantly reduced total walking distance and average walking velocity in the open field test. In a previous report, total distance of mouse movement was significantly reduced following chronic constriction injury (Blum, #9) of sciatic nerve-induced neuropathic pain, but was not reduced in a Complete Freund’s Adjuvant (CFA) model of inflammation, or in a spared nerve injury model (Urban et al., 2011). In contrast, significant differences in spontaneous foot lifting were detected in a CFA rat model, but not in a CCI mouse model (Djouhri et al., 2006; Mogil et al., 2010). Thus, as measures of spontaneous pain, spontaneous foot lifting does not necessarily correlate with open field test results. However, in our postoperative pain model, spontaneous foot lifting results were consistent with open field test results.

In our previous study, we used Pirt-GCaMP3 mice to monitor *in vivo* Ca^2+^ changes in primary sensory neurons in DRG, especially Ca^2+^ elevation in response to hindpaw stimuli. In the present study, we determined that spontaneous Ca^2+^ activity of primary sensory neurons in *in vivo* DRG Ca^2+^ imaging occurred under conditions of no stimulation to the hindpaw in the postoperative pain model. Intraplantar capsaicin injection prior to the incision abolished enhanced spontaneous Ca^2+^ activity of DRG primary sensory neurons, and significantly decreased both the number of Ca^2+^ response cells and Ca^2+^ transients of each soma caused by plantar stimuli. We conclude that spontaneous Ca^2+^ activity of DRG primary sensory neurons contributes to spontaneous pain, because intraplantar capsaicin injection significantly reduced both spontaneous pain and spontaneous Ca^2+^ activity of DRG primary sensory neurons. Our previous study suggested that IGF-2 expression in DRG was increased by plantar incision, but the increased IGF-2 expression was not detected with intraplantar capsaicin injection prior to the incision surgery (Tran et al., 2020). We interpret this to mean that the increase in IGF-2 resulting from the planar incision would cause so called “spontaneous” Ca^2+^ activity of DRG primary sensory neurons as an endogenous mediator. IGF-1 and IGF-2 can cross-activate and directly activate both IGF-1 and IGF-2 receptors (Rowzee et al., 2008). Each IGF receptor has different properties in physiological functions (Gary-Bobo et al., 2007; Hakuno and Takahashi, 2018). Further experiments will be needed to elucidate the mechanisms related to IGF signaling underlying post-operative pain.

In our postoperative pain model, the number of activated cells was significantly increased by stimulation with large brush, 100 g press, 300 g press, 50°C water, or 100 mM KCl, although not by acetone, compared to naïve animals and postoperative pain animals treated with intraplantar capsaicin injection. The number of cells activated by a mild stimulus like large brush was much less than numbers activated by harsher stimuli such as 100 g press, 300 g press, and 50 °C water. Additionally, we monitored only the surface of DRG using confocal microscopy, and it is possible that the actual cell number activated by mild stimuli like large brush would be greater if we could image deeper into the DRG. Other groups and our own results also showed that the activated cell number is different depending on the stimulus in *in vivo* DRG and trigeminal ganglia Ca^2+^ imaging (Kim et al., 2016; Chisholm et al., 2018; Leijon et al., 2019). More detailed analysis reveals that the increased cell number following plantar incision is mainly due to an increase in small to medium diameter cells. These results suggest that plantar incision has specific influences on primary sensory neuron subtypes. The present study shows that TRPV1-positive primary sensory neurons are involved in postoperatively enhanced Ca^2+^ responses. However, experiments with capsaicin injection did not indicate which specific primary sensory neuron subtype was involved in each enhanced postoperative pain behavior in our post-surgical pain model, although a plausible conclusion is that TRPV1 channel-positive neurons are key players in postoperative pain. Other researchers found that primary sensory neurons from TRPV1-cre animals labeled small diameter C fibers and a subset of medium Aδ fibers that innervate lamina I and II in the spinal cord, especially peptidergic primary sensory neurons. Our present and previous results are consistent with these findings (Bär et al., 2004; Cavanaugh et al., 2011; Mishra et al., 2011; Pogorzala et al., 2013; Le Pichon and Chesler, 2014). Additionally, Mitchell et al found that TRPV1-expressing Aδ fibers are involved in inflammatory pain which is controlled by resiniferatoxin, a strong TRPV1 channel agonist (Mitchell et al., 2010; Mitchell et al., 2014). Our next aim to understand further mechanisms underlying postoperative pain is to elucidate details of the role played by specific primary sensory neuron subtypes in each enhanced pain behavior using genetic CGRP-cre or MrgprD-cre mouse lines for peptidergic or non-peptidergic fiber positive primary sensory neurons, respectively (Le Pichon and Chesler, 2014). Previous studies showed that inhibition of CGRPα using CGRP_8-37_ alleviated mechanical and thermal pain postoperatively (Cowie et al., 2018). In addition, the decrease in skin temperature produced by acetone can reach to around 15-21°C. The threshold temperature for activation of transient receptor potential melastatin 8 (TRPM8) and transient receptor potential ankyrin 1 (TRPA1) channels is around 25°C and 17°C, respectively (Dhaka et al., 2006; Chen, 2015). Involvement of TRPA1 channels in spontaneous and mechanical pain in a postoperative pain model has been reported (Wei et al., 2012). Also, acetone-induced cooling stimulation in the present study could activate TRPM8 but possibly not TRPA1 channels. Further experiments are needed to confirm the involvement of TRPA1 channel in postoperative pain including cold-induced pain.

Some studies showed neuronal plasticity in spinal cord involving increased expression of c-Fos in postoperative pain (Zhu et al., 2006; Guo et al., 2014; Gu et al., 2019; Xu et al., 2019). Only a few studies, using Ca^2+^ dye or genetically-encoded calcium indicators, have examined the contribution to postoperative pain of Ca^2+^ activity and Ca^2+^ signaling in spinal cord. We tried to monitor Ca^2+^ activity and Ca^2+^ signaling at primary sensory neuron central terminals in spinal cord using the Pirt-GCaMP3 mouse model. In each spinal cord slice, we used clearly visible green fluorescence to identify primary sensory nerve fibers in the area of lamina I and II. Corresponding to *in vivo* DRG Ca^2+^ imaging, Ca^2+^ transients and elevation induced by capsaicin and high KCl (50 mM) in the ipsilateral side were significantly enhanced compared to activity in the contralateral side. The findings suggest that plantar incision induces plasticity-related changes in primary sensory neuron central terminals in spinal cord.

In summary, in a postoperative pain model, primary sensory neuronal activity is dramatically enhanced, and subcutaneous capsaicin injection prior to incision surgery causes TRPV1-expressing neuronal nerve fiber degeneration followed by the alleviation of spontaneous, mechanical, and thermal pain, and also significantly reduces Ca^2+^ elevation and activity of DRG primary sensory neurons. In a previous clinical study, usefulness of a capsaicin patch for postoperative pain was reported (Tamburini et al., 2018). Although capsaicin elevates Ca^2+^ concentration through the activation of TRPV1 channels, the capsaicin-mediated Ca^2+^ elevation is weaker than resiniferatoxin-mediated Ca^2+^ elevation, but strong enough to control TRPV1-mediated pain conditions; resiniferatoxin apparently can control these various TRPV1-related pain types including postoperative pain (Szallasi and Blumberg, 1990; Neubert et al., 2003; Karai et al., 2004; Neubert et al., 2008; Hockman et al., 2018; Raithel et al., 2018). Our findings support the use of vanilloid agonist or TRPV1 agonist treatment options for postoperative pain.

## Acknowledgement

This work was supported by National Institutes of Health Grant (R01DE026677 to Y.K.), UTHSA startup fund (to Y.K.) and a STAR Award (to Y.K.) from University of Texas system.

## References

Ahmad S, De Oliveira GS, Jr., Bialek JM, McCarthy RJ (2014) Thermal quantitative sensory testing to predict postoperative pain outcomes following gynecologic surgery. Pain Med 15:857–864.

Apfelbaum JL, Chen C, Mehta SS, Gan TJ (2003) Postoperative pain experience: results from a national survey suggest postoperative pain continues to be undermanaged. Anesth Analg 97:534–540, table of contents.

Banik RK, Subieta AR, Wu C, Brennan TJ (2005) Increased nerve growth factor after rat plantar incision contributes to guarding behavior and heat hyperalgesia. Pain 117:68–76.

Bär KJ, Schaible HG, Bräuer R, Halbhuber KJ, von Banchet GS (2004) The proportion of TRPV1 protein-positive lumbar DRG neurones does not increase in the course of acute and chronic antigen-induced arthritis in the knee joint of the rat. Neurosci Lett 361:172–175.

Benyamin R, Trescot AM, Datta S, Buenaventura R, Adlaka R, Sehgal N, Glaser SE, Vallejo R (2008) Opioid complications and side effects. Pain Physician 11:S105–120.

Binzen U, Greffrath W, Hennessy S, Bausen M, Saaler-Reinhardt S, Treede RD (2006) Co-expression of the voltage-gated potassium channel Kv1.4 with transient receptor potential channels (TRPV1 and TRPV2) and the cannabinoid receptor CB1 in rat dorsal root ganglion neurons. Neuroscience 142:527–539.

Carroll I, Barelka P, Wang CK, Wang BM, Gillespie MJ, McCue R, Younger JW, Trafton J, Humphreys K, Goodman SB, Dirbas F, Whyte RI, Donington JS, Cannon WB, Mackey SC (2012) A pilot cohort study of the determinants of longitudinal opioid use after surgery. Anesth Analg 115:694–702.

Cavanaugh DJ, Chesler AT, Bráz JM, Shah NM, Julius D, Basbaum AI (2011) Restriction of transient receptor potential vanilloid-1 to the peptidergic subset of primary afferent neurons follows its developmental downregulation in nonpeptidergic neurons. J Neurosci 31:10119–10127.

Chang H, Wang Y, Wu H, Nathans J (2014) Flat mount imaging of mouse skin and its application to the analysis of hair follicle patterning and sensory axon morphology. J Vis Exp:e51749.

Chaplan SR, Bach FW, Pogrel JW, Chung JM, Yaksh TL (1994) Quantitative assessment of tactile allodynia in the rat paw. J Neurosci Methods 53:55–63.

Chen J (2015) The evolutionary divergence of TRPA1 channel: heat-sensitive, cold-sensitive and temperature-insensitive. Temperature (Austin) 2:158–159.

Chisholm KI, Khovanov N, Lopes DM, La Russa F, McMahon SB (2018) Large Scale In Vivo Recording of Sensory Neuron Activity with GCaMP6. eNeuro 5.

Cowie AM, Moehring F, O’Hara C, Stucky CL (2018) Optogenetic Inhibition of CGRPα Sensory Neurons Reveals Their Distinct Roles in Neuropathic and Incisional Pain. J Neurosci 38:5807–5825.

Dhaka A, Viswanath V, Patapoutian A (2006) Trp ion channels and temperature sensation. Annu Rev Neurosci 29:135–161.

Djouhri L, Koutsikou S, Fang X, McMullan S, Lawson SN (2006) Spontaneous pain, both neuropathic and inflammatory, is related to frequency of spontaneous firing in intact C-fiber nociceptors. J Neurosci 26:1281–1292.

Dráberová E, Lukás Z, Ivanyi D, Viklický V, Dráber P (1998) Expression of class III beta-tubulin in normal and neoplastic human tissues. Histochem Cell Biol 109:231–239.

Farahani RM, Rezaei-Lotfi S, Simonian M, Xaymardan M, Hunter N (2019) Neural microvascular pericytes contribute to human adult neurogenesis. J Comp Neurol 527:780–796.

Gary-Bobo M, Nirdé P, Jeanjean A, Morère A, Garcia M (2007) Mannose 6-phosphate receptor targeting and its applications in human diseases. Curr Med Chem 14:2945–2953.

Green DP, Limjunyawong N, Gour N, Pundir P, Dong X (2019) A Mast-Cell-Specific Receptor Mediates Neurogenic Inflammation and Pain. Neuron 101:412–420.e413.

Gu HW, Xing F, Jiang MJ, Wang Y, Bai L, Zhang J, Li TT, Zhang W, Xu JT (2019) Upregulation of matrix metalloproteinase-9/2 in the wounded tissue, dorsal root ganglia, and spinal cord is involved in the development of postoperative pain. Brain Res 1718:64–74.

Gulyás M, Bencsik N, Pusztai S, Liliom H, Schlett K (2016) AnimalTracker: An ImageJ-Based Tracking API to Create a Customized Behaviour Analyser Program. Neuroinformatics 14:479–481.

Guo R, Zhao Y, Zhang M, Wang Y, Shi R, Liu Y, Xu J, Wu A, Yue Y, Wu J, Guan Y, Wang Y (2014) Down-regulation of Stargazin inhibits the enhanced surface delivery of α-amino-3-hydroxy-5-methyl-4-isoxazole propionate receptor GluR1 subunit in rat dorsal horn and ameliorates postoperative pain. Anesthesiology 121:609–619.

Hakuno F, Takahashi SI (2018) IGF1 receptor signaling pathways. J Mol Endocrinol 61:T69–t86.

Hockman TM, Cisternas AF, Jones B, Butt MT, Osborn KG, Steinauer JJ, Malkmus SA, Yaksh TL (2018) Target engagement and histopathology of neuraxial resiniferatoxin in dog. Vet Anaesth Analg 45:212–226.

Hsieh YL, Lin CL, Chiang H, Fu YS, Lue JH, Hsieh ST (2012) Role of peptidergic nerve terminals in the skin: reversal of thermal sensation by calcitonin gene-related peptide in TRPV1-depleted neuropathy. PLoS One 7:e50805.

Joksimovic SL, Joksimovic SM, Tesic V, García-Caballero A, Feseha S, Zamponi GW, Jevtovic-Todorovic V, Todorovic SM (2018) Selective inhibition of Ca(V)3.2 channels reverses hyperexcitability of peripheral nociceptors and alleviates postsurgical pain. Sci Signal 11.

Kang S, Wu C, Banik RK, Brennan TJ (2010) Effect of capsaicin treatment on nociceptors in rat glabrous skin one day after plantar incision. Pain 148:128–140.

Karai L, Brown DC, Mannes AJ, Connelly ST, Brown J, Gandal M, Wellisch OM, Neubert JK, Olah Z, Iadarola MJ (2004) Deletion of vanilloid receptor 1-expressing primary afferent neurons for pain control. J Clin Invest 113:1344–1352.

Katsetos CD, Frankfurter A, Christakos S, Mancall EL, Vlachos IN, Urich H (1993) Differential localization of class III, beta-tubulin isotype and calbindin-D28k defines distinct neuronal types in the developing human cerebellar cortex. J Neuropathol Exp Neurol 52:655–666.

Kawamata M, Watanabe H, Nishikawa K, Takahashi T, Kozuka Y, Kawamata T, Omote K, Namiki A (2002) Different mechanisms of development and maintenance of experimental incision-induced hyperalgesia in human skin. Anesthesiology 97:550–559.

Kim YS, Chu Y, Han L, Li M, Li Z, LaVinka PC, Sun S, Tang Z, Park K, Caterina MJ, Ren K, Dubner R, Wei F, Dong X (2014) Central terminal sensitization of TRPV1 by descending serotonergic facilitation modulates chronic pain. Neuron 81:873–887.

Kim YS, Anderson M, Park K, Zheng Q, Agarwal A, Gong C, Saijilafu Young L, He S, LaVinka PC, Zhou F, Bergles D, Hanani M, Guan Y, Spray DC, Dong X (2016) Coupled Activation of Primary Sensory Neurons Contributes to Chronic Pain. Neuron 91:1085–1096.

Le Pichon CE, Chesler AT (2014) The functional and anatomical dissection of somatosensory subpopulations using mouse genetics. Front Neuroanat 8:21.

Leijon SCM, Neves AF, Breza JM, Simon SA, Chaudhari N, Roper SD (2019) Oral thermosensing by murine trigeminal neurons: modulation by capsaicin, menthol and mustard oil. J Physiol 597:2045–2061.

Li F, Yang W, Jiang H, Guo C, Huang AJW, Hu H, Liu Q (2019) TRPV1 activity and substance P release are required for corneal cold nociception. Nat Commun 10:5678.

Lovich-Sapola J, Smith CE, Brandt CP (2015) Postoperative pain control. Surg Clin North Am 95:301–318.

Macrae WA (2008) Chronic post-surgical pain: 10 years on. Br J Anaesth 101:77–86.

Mishra SK, Tisel SM, Orestes P, Bhangoo SK, Hoon MA (2011) TRPV1-lineage neurons are required for thermal sensation. Embo j 30:582–593.

Mitchell K, Lebovitz EE, Keller JM, Mannes AJ, Nemenov MI, Iadarola MJ (2014) Nociception and inflammatory hyperalgesia evaluated in rodents using infrared laser stimulation after Trpv1 gene knockout or resiniferatoxin lesion. Pain 155:733–745.

Mitchell K, Bates BD, Keller JM, Lopez M, Scholl L, Navarro J, Madian N, Haspel G, Nemenov MI, Iadarola MJ (2010) Ablation of rat TRPV1-expressing Adelta/C-fibers with resiniferatoxin: analysis of withdrawal behaviors, recovery of function and molecular correlates. Mol Pain 6:94.

Mogil JS, Graham AC, Ritchie J, Hughes SF, Austin JS, Schorscher-Petcu A, Langford DJ, Bennett GJ (2010) Hypolocomotion, asymmetrically directed behaviors (licking, lifting, flinching, and shaking) and dynamic weight bearing (gait) changes are not measures of neuropathic pain in mice. Mol Pain 6:34.

Neubert JK, Karai L, Jun JH, Kim HS, Olah Z, Iadarola MJ (2003) Peripherally induced resiniferatoxin analgesia. Pain 104:219–228.

Neubert JK, Mannes AJ, Karai LJ, Jenkins AC, Zawatski L, Abu-Asab M, Iadarola MJ (2008) Perineural resiniferatoxin selectively inhibits inflammatory hyperalgesia. Mol Pain 4:3.

Pogorzala LA, Mishra SK, Hoon MA (2013) The cellular code for mammalian thermosensation. J Neurosci 33:5533–5541.

Raithel SJ, Sapio MR, LaPaglia DM, Iadarola MJ, Mannes AJ (2018) Transcriptional Changes in Dorsal Spinal Cord Persist after Surgical Incision Despite Preemptive Analgesia with Peripheral Resiniferatoxin. Anesthesiology 128:620–635.

Roh J, Hwang SM, Lee SH, Lee K, Kim YH, Park CK (2020) Functional Expression of Piezo1 in Dorsal Root Ganglion (DRG) Neurons. Int J Mol Sci 21.

Rowzee AM, Lazzarino DA, Rota L, Sun Z, Wood TL (2008) IGF ligand and receptor regulation of mammary development. J Mammary Gland Biol Neoplasia 13:361–370.

Schreml S, Szeimies RM, Karrer S, Heinlin J, Landthaler M, Babilas P (2010) The impact of the pH value on skin integrity and cutaneous wound healing. J Eur Acad Dermatol Venereol 24:373–378.

Szallasi A, Blumberg PM (1990) Resiniferatoxin and its analogs provide novel insights into the pharmacology of the vanilloid (capsaicin) receptor. Life Sci 47:1399–1408.

Tamburini N, Bollini G, Volta CA, Cavallesco G, Maniscalco P, Spadaro S, Qurantotto F, Ragazzi R (2018) Capsaicin patch for persistent postoperative pain after thoracoscopic surgery, report of two cases. J Vis Surg 4:51.

Thapa P, Euasobhon P (2018) Chronic postsurgical pain: current evidence for prevention and management. Korean J Pain 31:155–173.

Tominaga M, Tominaga T (2005) Structure and function of TRPV1. Pflugers Arch 451:143–150.

Tran PV, Johns ME, McAdams B, Abrahante JE, Simone DA, Banik RK (2020) Global transcriptome analysis of rat dorsal root ganglia to identify molecular pathways involved in incisional pain. Mol Pain 16:1744806920956480.

Uhelski ML, McAdams B, Johns ME, Kabadi RA, Simone DA, Banik RK (2020) Lack of relationship between epidermal denervation by capsaicin and incisional pain behaviours: A laser scanning confocal microscopy study in rats. Eur J Pain 24:1197–1208.

Urban R, Scherrer G, Goulding EH, Tecott LH, Basbaum AI (2011) Behavioral indices of ongoing pain are largely unchanged in male mice with tissue or nerve injury-induced mechanical hypersensitivity. Pain 152:990–1000.

Wang YX, Pettus M, Gao D, Phillips C, Scott Bowersox S (2000) Effects of intrathecal administration of ziconotide, a selective neuronal N-type calcium channel blocker, on mechanical allodynia and heat hyperalgesia in a rat model of postoperative pain. Pain 84:151–158.

Wei H, Karimaa M, Korjamo T, Koivisto A, Pertovaara A (2012) Transient receptor potential ankyrin 1 ion channel contributes to guarding pain and mechanical hypersensitivity in a rat model of postoperative pain. Anesthesiology 117:137–148.

Wu H, Williams J, Nathans J (2012) Morphologic diversity of cutaneous sensory afferents revealed by genetically directed sparse labeling. Elife 1:e00181.

Xu B, Mo C, Lv C, Liu S, Li J, Chen J, Wei Y, An H, Ma L, Guan X (2019) Post-surgical inhibition of phosphatidylinositol 3-kinase attenuates the plantar incision-induced postoperative pain behavior via spinal Akt activation in male mice. BMC Neurosci 20:36.

Zhu CZ, Nikkel AL, Martino B, Bitner RS, Decker MW, Honore P (2006) Dissociation between post-surgical pain behaviors and spinal Fos-like immunoreactivity in the rat. Eur J Pharmacol 531:108–117.

